# Analysis of a hypomorphic *mei-P26* mutation reveals coordination between developmental programming of germ cells and meiotic chromosome dynamics

**DOI:** 10.1101/2024.11.22.624911

**Authors:** Joseph Terry, Ally Solomon, Amanda M. Powell, Patrick Terry, Oscar Bautista, Marty Landes, Erica Berent, Elizabeth T. Ables, Nicole Crown

## Abstract

Female gametogenesis in *Drosophila* requires differentiation and mitotic division of germ cells, acquisition of oocyte fate, and entry into meiosis. Each of these processes is well understood individually; however, little is known about the mechanisms that ensure proper temporal integration of germ cell differentiation and meiotic chromosome dynamics. Here, we take advantage of a hypomorphic mutation in *mei-P26*, a well-characterized gene with multiple diverse functions in germ cell development, to determine the consequences of disrupting the coordination between development and meiosis. While null mutations in *mei-P26* lead to tumorous ovaries, the hypomorphic allele *mei-P26*^*1*^ allows sufficient germ cell differentiation and fertility to support analysis of meiotic chromosome dynamics. Unlike wildtype germaria, 60% of cysts in *mei-P26*^*1*^ germaria co-express the differentiation factor Bag of marbles (Bam) and the oocyte specification factor Orb, suggesting that mitotic division is delayed. In this context, the synaptonemal complex rarely assembles into full length continuous tracks and instead is missing or present only as foci. Despite these phenotypes, meiotic double-strand breaks still form and are repaired as crossovers, but the crossovers are mis-patterned and form in centromere proximal regions rather than chromosome arms. The strength of crossover interference is significantly reduced and the centromere effect is lost, but crossover assurance is intact and the meiosis-specific machinery is used to form crossovers. We suggest a model where the failure to exit mitosis in a timely fashion causes cells to enter meiosis while still receiving mitotic signals, resulting in abnormal meiotic chromosome dynamics and impaired crossover formation.

**Article summary:** Female gametogenesis in *Drosophila* requires the precise temporal integration of germ cell differentiation and meiotic entry. Using a hypomorphic mutation in *mei-P26*, we investigated how disrupting this coordination affects meiotic chromosome dynamics. In *mei-P26*^*1*^ mutants, delayed mitotic exit causes cells to enter meiosis while still receiving mitotic signals. This results in fragmented synaptonemal complex and the loss of some but not all CO patterning mechanisms. These results suggest that timely mitotic exit is critical for establishing the proper regulatory landscape required for meiotic recombination.

## Introduction

Germ cells are unipotent and essential to the formation of all cells in the next generation, but they must also undergo a unique cell cycle program – meiosis – that uses specialized chromosome dynamics to go from a diploid to a haploid genome. For germ cells that arise from a germline-restricted stem cell population, initiation of the meiotic cell cycle is tightly coordinated with acquisition of the oocyte or sperm fate to result in viable gametes. This is well-illustrated in the female *Drosophila* germline, where an ovarian germline stem cell (GSC) produces a committed oocyte progenitor (called the cystoblast) that undergoes four mitotic divisions while simultaneously initiating the chromosome dynamics of meiosis (Figure 1, [1]). Genetic analysis of mutants defective in stem cell differentiation has provided a deep understanding about the developmental control of germline stem cell differentiation and oocyte specification. Additionally, the timing of meiosis relative to GSC differentiation and oocyte specification is well known. Much less is known, however, about the mechanism that integrates the timing of meiotic chromosome dynamics (such as chromosome pairing and recombination) with developmental programming, and how disrupting this integration might affect meiosis.

**Figure 1.**
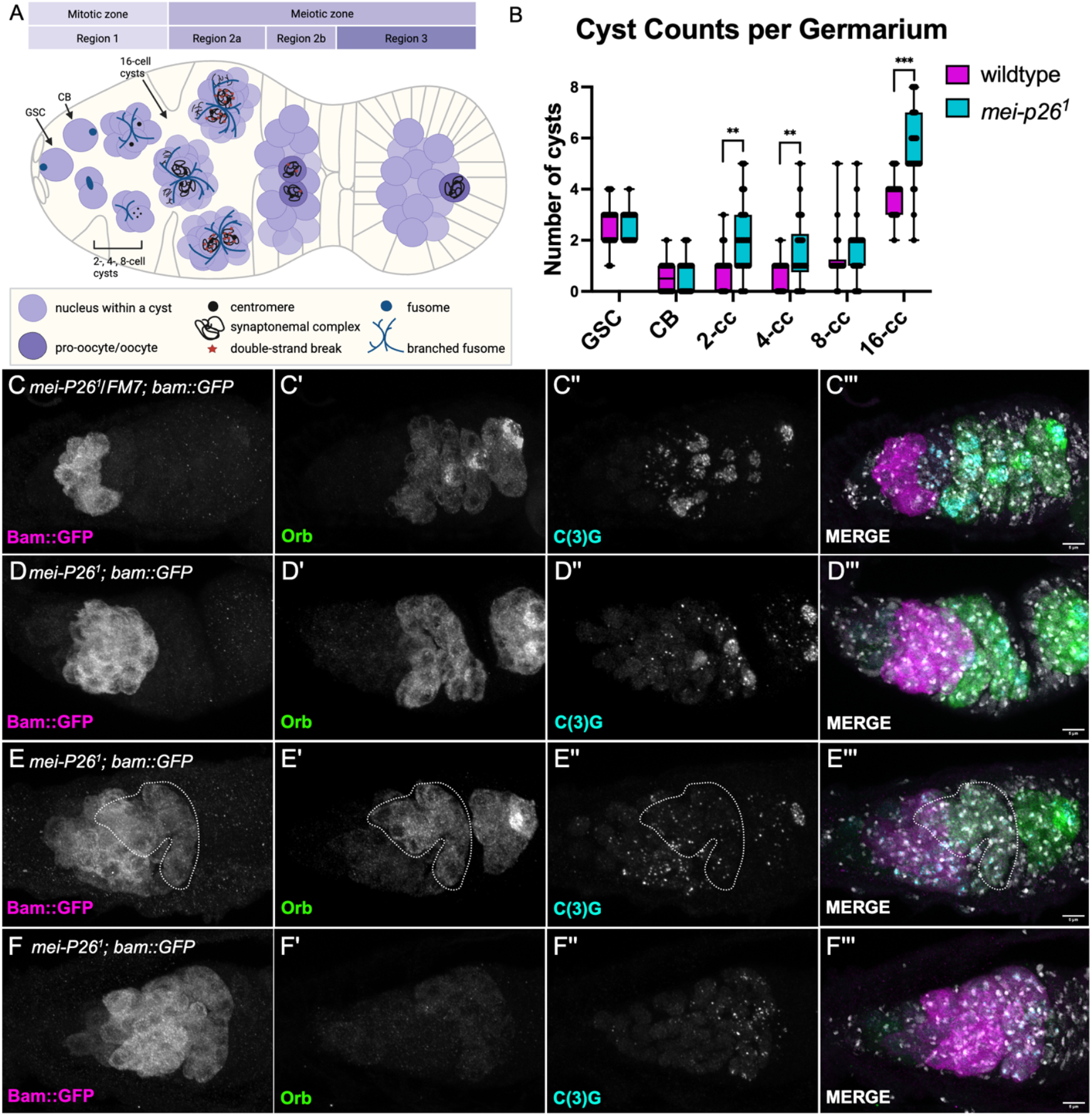
**A)** Diagram of the Drosophila germarium. **B)** There are significantly more cysts at the 2-, 4-, and 16-cell stages in *mei-P26*^*1*^ compared to wildtype (see text for details, p<0.001). **C-C’’’)** In heterozygote controls, Bam and Orb are never co-expressed (n = 20 germaria). **D-D’’’)** 24% of *mei-P26*^*1*^ germaria express Bam and Orb in mutually exclusive patterns similar to *mei-P26*^*1*^ heterozygotes (n = 29 germaria). **E-E’’’)** In *mei-P26*^*1*^ homozygotes, Bam and Orb are co-expressed in at least one cyst in 62% of germaria (n = 29 germaria). The region of overlap is drawn with a dashed line. **F-F’’’)** In 14% of *mei-P26*^*1*^ germaria, Bam is expressed but Orb is not (n = 29 germaria). **C’’’-F’’’)** DAPI staining in gray.

GSC identity is maintained by signals from adjacent somatic cells at the anterior tip of the germarium (Figure 1) [1]. Upon GSC division, the cystoblast is displaced from these signals, triggering activation of differentiation factors such as Bag of marbles (Bam). The cystoblast undergoes exactly four rounds of mitosis with incomplete cytokinesis, producing interconnected 2-, 4-, 8-, and 16-cell cysts linked by a membranous organelle called the fusome. As cysts divide, they move posteriorly from region 1 into region 2a, and all cells of the cyst accumulate the oocyte specification factor Orb. In region 2b, Orb hyperaccumulates in two cells, termed pro-oocytes. By region 3, Orb is restricted to a single cell, specifying the oocyte, while the remaining 15 cells differentiate as nurse cells (reviewed in [1]).

At the same time germ cells are undergoing differentiation and oocyte specification, the chromosome dynamics of meiosis occur. Homologous chromosomes pair and build the synaptonemal complex, a tripartite zipper-like structure that facilitates homolog interactions. Next, programmed DNA double-strand breaks are made and repaired using homologous recombination to form crossovers (COs). COs are stabilized into chiasma by sister chromatid cohesion and are essential for correct chromosome segregation, as failure to form or correctly position COs leads to chromosome nondisjunction and aneuploid gametes [2, 3]. CO formation and placement are regulated by three conserved patterning mechanisms: crossover assurance, which ensures at least one CO per homolog pair; CO interference, which spaces multiple COs farther apart than expected by chance; and the centromere effect, which suppresses COs near centromeres. Collectively, these mechanisms result in a stereotyped CO distribution, such that *Drosophila* females have one to two COs per chromosome arm, primarily in the medial portion of the arms with few COs near centromeres or telomeres [4].

Each hallmark event of meiotic prophase is precisely coordinated with germ cell differentiation and oocyte fate acquisition (Figure 1). Homologous chromosome pairing begins in the 8-cell cyst, when synaptonemal complex proteins are loaded onto centromeres to facilitate centromere clustering and homologous chromosome pairing (how chromosomes pair prior to a final round of mitosis is unknown [5, 6]). As 16-cell cysts progress posteriorly, the synaptonemal complex is loaded at five to six sites on chromosome arms [5, 7]. Upon entry into region 2a, the synaptonemal complex is fully loaded onto chromosome arms in continuous tracks, and double-strand breaks are made by the Spo-11 ortholog Mei-W68 [8, 9]. DSB repair is completed by region 3 and Vilya foci, a cytological marker of COs, appear in region 2b [10]. By region 3, meiotic recombination and CO formation are complete [1, 4].

Despite our rich knowledge of GSC differentiation, oocyte specification, meiotic chromosome dynamics, and the developmental timing of meiosis, the mechanisms that integrate these processes remain largely unknown. This knowledge gap exists primarily because strong differentiation mutants cause tumorous ovaries or sterility, limiting analysis of meiotic progression. To address this problem, we used a hypomorphic allele of the well-characterized gene *mei-P26* that supports sufficient germline development to maintain fertility. *mei-P26* encodes a TRIM-NHL protein required for GSC differentiation in the ovary and testes [11]. The protein functions molecularly to repress *nanos* translation in the cystoblast via Bam, Bgcn, and Sxl [12], promote differentiation by inhibiting the miRNA pathway [13], and increase the rate of protein translation during cyst divisions by upregulating Target of Rapamycin (Tor) activity [14]. While null alleles of *mei-P26* result in tumorous ovaries and testes [13], previous studies suggested that germ cells carrying the hypomorphic allele (*mei-P26*^*1*^) complete cyst development but have meiotic defects, including X chromosome nondisjunction and reduced CO frequency [11]. Additionally, synaptonemal complex assembly appears incomplete in *mei-P26*^*1*^ ovaries [15], suggesting a role for Mei-P26 in meiosis beyond differentiation.

Here, we show that germ cells in *mei-P26*^*1*^ ovaries initially receive both mitotic and oocyte specification signals at the same time. Germline cysts accumulate irregularly, and over half of germaria contain cysts that improperly co-express Bam and Orb. Although germ cells in *mei-P26*^*1*^ mutants enter meiosis, few nuclei assemble continuous, full-length synaptonemal complexes. Despite the failure to make robust synaptonemal complexes, double-stranded breaks and COs still form, but COs fail to be properly patterned. We propose that ovarian germ cells must transition from a mitotic to a meiotic cell identity in the correct temporal sequence in order for meiotic chromosome dynamics to occur normally.

## Materials and Methods

### Drosophila husbandry

All *Drosophila* stocks were maintained at 25 degrees Celsius on standard cornmeal media. *yw P{w[+mC]=lacW mei-P26*^*1*^/C(1)DX was obtained from Scott Hawley, and the *y mei-9*^*a*^/FM7w stock and *mei-P22*^*P22*^ stock from Jeff Sekelsky. The *Bam::GFP* stock is *y[1] w[*];; bam-2XTY1-SGFP-V5-preTEVBLRP-3XFLAG[VK00033]* (VDRC line #318001). Multiple different stocks were used as controls over the course these experiments. For immunostaining, either *y*^*1*^ or *w*^*1118*^ were used as controls. For the *Bam::GFP* fusion, *mei-P26*^*1*^/FM7;; *Bam::GFP* siblings were used as controls. For crossover scoring, Oregon-RM or *y*^*1*^ *w*^*1118*^ were used as controls.

### Immunostaining

2-3 day old mated females were put on yeast paste overnight. Ovaries were dissected in PBS and fixed for 20 minutes in 1000 μl of solution containing 2% paraformaldehyde (Ted Pella cat. no. 18505), 0.5% IGEPAL CA-630 (Thermo Fisher cat. no. 18896)), 200 μl PBS and 600 μl heptane. Ovaries were then washed three times for ten minutes each in PBS with 0.1% Tween-20 (PBST), blocked for one hour at room temperature in PBS with 1% BSA (MP Biomedicals cat. no. 152401) and incubated with primary antibody diluted in PBST overnight at 4 degrees. Ovaries were then washed three times in PBST and incubated in secondary antibody diluted in PBST for 4 hours at room temperature. DAPI was added for the last 10 minutes at a concentration of 1 μl /ml. Ovaries were washed again three times for 15 minutes each in PBST. All wash steps and antibody incubations were done while nutating. Ovaries were mounted in ProLong Glass (Invitrogen cat. no. P36980) and allowed to cure for the manufacturer’s suggested time.

The following primary antibodies were used: mouse anti-phospho-H2AV (1:500) [16]; mouse anti-C(3)G 1A8-1G2 (1:500) [17]; guinea pig anti-CENP-C (1:500), gift from Kim McKim [18]; mouse anti-Hts (1:500), clone 1B1, Developmental Studies Hybridoma Bank; mouse anti-Cyclin A (1:10), clone A12, Developmental Studies Hybridoma Bank; mouse anti-Orb (1:20 or 1:250), clone 6H, Developmental Studies Hybridoma Bank; chicken anti-GFP (1:2000) AB13970, ABCAM.

The following secondary antibodies were used at 1:500: anti-IgG1 Alexa488 (Thermo Fisher A21121); anti-IgG2b Alexa594 (Thermo Fisher A21145); anti-mouse Alexa488 (Thermo Fisher A11001); anti-mouse Alexa594 (Thermo Fisher A11005); anti-guinea pig Alexa647 (Thermo Fisher A21450); anti-rabbit Alexa647 (Thermo Fisher A31573); anti-chicken Alexa488 (Thermo Fisher A11039).

Ovaries were imaged on a Leica Stellaris 5 confocal microscope using an HC PL APO 63x/1.4 NA Oil objective. Images were deconvolved using the Leica Lightning. Bam-GFP images were acquired on a Zeiss LSM 700 confocal microscope using a 63x/1.4 NA oil objective.

### Quantification of γH2AV and C(3)G fluorescence intensity

Confocal z-stacks were processed and analyzed in FIJI/ImageJ v1.54p. Individual cysts were segmented in three dimensions by manually delineating regions of interest (ROIs), after which ROIs were visually inspected and manually corrected where necessary. For each cyst and channel, raw integrated density and area were measured on every segmented z-slice. Per-cyst mean fluorescence intensity was calculated as the sum of raw integrated density across all segmented slices divided by the sum of segmented area across slices, yielding a single area-weighted stack mean for each cyst. The same ROI set was used to quantify both *γ*H2Av and C(3)G signals γH2AV signal is present in non-meiotic cells of the germarium, so analysis was limited to *γ*H2AV signal that overlapped with C(3)G using a masking approach. Semi-automated segmentation was performed with Labkit v0.1.18. A pixel-classification model was trained on representative images spanning the range of observed synaptonemal complex morphologies (foci, discontinuous, and continuous) and applied to generate binary masks. Predicted masks were reviewed manually and corrected as needed before conversion to ROIs for downstream quantification. No background subtraction, intensity rescaling, bleaching correction, or other preprocessing was applied before measurement; all fluorescence measurements were made from raw image data. Images included in a given quantitative comparison were acquired using identical imaging settings.

### Statistical analysis – mixed effect models

Because our data is structured (genotype → cysts within the same germarium → individual cyst), some data sets are unbalanced, and some data sets are not normally distributed,we used mixed effects models to simultaneously account for multiple predictors in one cohesive framework.

All statistical analyses were performed in R v4.4.3. Cyst-level models included Genotype, Stage (GSC, CB, 2-, 4-, 8-, and 16-cell), and their interaction as fixed effects, with ImageID (Germarium) included as a random intercept to account for clustering of cysts within images. Categorical predictors were coded using sum-to-zero contrasts for Type III inference. Fixed effects were assessed using Type III Wald χ^2^ tests with car::Anova, pairwise comparisons were obtained from estimated marginal means using emmeans with Tukey-adjusted p-values, and model diagnostics were assessed using DHARMa and performance. Exact p-values are reported for p≥0.001; smaller values are reported as p<0.001.

Cyst-composition counts were analyzed at the germarium level using a Conway-Maxwell Poisson mixed model implemented in glmmTMB. Candidate count distributions were compared using AIC and residual diagnostics on the same dataset. Poisson and negative-binomial models showed systematic lack of fit in DHARMa diagnostics, indicating misspecification of the mean– variance relationship. The final model therefore used a Conway–Maxwell–Poisson distribution,which allows flexible modeling of both under- and overdispersion. Model fit was evaluated using DHARMa, and explanatory power was summarized using likelihood-ratio R2 (Cox–Snell) from MuMIn::r.squaredLR, computed relative to an intercept-only COM-Poisson model.

H2Av fluorescence intensity was analyzed as log-transformed per-cyst mean intensity using a hetergeneous variance mixed model implemented in glmmTMB. The number of segmented z-slices was evaluated as a nuisance covariate and omitted from the final model; it was not used to divide the already averaged intensity measure. Preliminary models assuming homoscedastic residual variance showed systematic violations of model assumptions based on DHARMa diagnostics, including heteroscedasticity across fitted values and predictor levels. To accommodate non-constant residual variance, a model with a dispersion component was fitted, allowing residual variance to vary as a function of Genotype and Stage. Explanatory power was summarized using a likelihood-ratio based pseudo-R^2^ (MuMIn::r.squaredLR) relative to an intercept-only model, as variance-partitioning R^2^ was not applicable under the dispersion-extended model structure. An analogous model replacing MStage with Region was fitted under the same specification for a 16-cell subset.

C3G fluorescence intensity was analyzed at the cyst level using a zero-inflated gamma mixed model implemented in glmmTMB. Initial Gaussian, log-Gaussian, Gamma, and Tweedie models showed persistent lack of fit under residual diagnostics, with structured deviations driven by a distinct low-intensity subpopulation predominantly observed in meiP26 cysts. Based on an empirically supported threshold in wild-type cysts, low-intensity observations were classified as non-expressing, enabling a two-process model formulation. C3G intensity was therefore modeled using a zero-inflated Gamma GLMM. The zero-inflation component captured the probability of low signals, while the conditional Gamma component modeled variation in C3G intensity among expressing cysts. Explanatory power was quantified using Nakagawa’s marginal and conditional R^2^ for mixed models.

C(3)G morphological phenotypes were analyzed using separate Bayesian-penalized binomial mixed models, with each model coding the focal phenotype as 1 and all other scored phenotypes as 0. Because complete or quasi-complete separation occurred in some Genotype × Stage combinations, models were fit using bglmer with zero-centered weakly informative priors on fixed effects ([19] v3.1). Continuous and discontinuous phenotypes were analyzed only in 8- and 16-cell cysts. Model performance was summarized using Nakagawa’s marginal and conditional R^2^ to quantify variance explained by fixed effects and the full mixed model structure. Of note, when cysts contained multiple synaptonemal complex morphologies, they were counted separately in each category, thus the percentages do not equal 100%.

All code can be found at https://github.com/josephterry/MeiP26_CrownLab.

### Statistical analysis - other

Statistical analyses were performed in Graphpad Prism version 10 (GraphPad Software, Boston, Massachusetts USA, www.graphpad.com).

### CO scoring

To score crossover frequencies and distributions, we used recessive marker scoring. For chromosome 2L, females that were *net dpp*^*d-ho*^ *dpy b pr cn*/*+* were crossed to homozygous *net dpp*^*d-ho*^ *dpy b pr* males. For chromosome 3, females that were *ru*^*1*^ *h*^*1*^ *Diap*^*th1*^ *st*^*1*^ *p*^*p*^ *cu*^*1*^ *sr*^*1*^ *e*^*s*^/+ were crossed to homozygous *ru*^*1*^ *h*^*1*^ *Diap*^*th1*^ *st*^*1*^ *p*^*p*^ *cu*^*1*^ *sr*^*1*^ *e*^*s*^ males. Crossovers were identified by switches from the mutant to wildtype phenotypes (and vice versa) for each gene in the offspring. For each mutant genotype, the recessive marker chromosome was crossed into the mutant background.

CO frequencies in Table 1 and Supplemental Table 1 are presented as centiMorgans (cM), which is (# of COs in an interval/total sample size)*100. Fisher’s exact test with a Bonferroni correction for multiple comparisons was used to determine statistical differences in CO frequencies.

Crossover frequencies on chromosome 2L (cM)
|  | *net-dpp<sup>d-ho</sup>* | *dpp<sup>d-ho</sup>-dpy* | *dpy-b* | *b-pr* | *pr-cn<sup>^</sup>* | total | n |
| --- | --- | --- | --- | --- | --- | --- | --- |
| **Oregon-RM** | **2.68**<br>(1.66-3.69) | **3.4**<br>(2.26-4.54) | **27.70**<br>(24.89-30.52) | **5.36**<br>(3.94-6.77) | **1.54**<br>(0.77-2.32) | **40.68**<br>(39.79-41.57) | 971 |
| ***mei-p26<sup>1</sup>*** | **0.22\*\***<br>(0.03-0.4) | **0.82\*\***<br>(0.45-1.18) | **8.95\*\***<br>(7.79-10.11) | **9.17\***<br>(7.99-10.34) | **24.74\*\***<br>(22.99-26.50) | **43.89**<br>(43.0-44.78) | 2321 |
| ***mei-9<sup>a</sup>*** | **0.52\***<br>(0.01-1.03) | **0.39\*\***<br>(-0.5-0.83) | **2.34\*\***<br>(1.27-3.4) | **0.52\*\***<br>(0.01- 1.03) | **0.13\***<br>(-0.12 – 0.38) | **3.90\*\***<br>(3.0-4.79) | 770 |
| ***mei-9<sup>a</sup> mei-p26<sup>1</sup>*** | **0** | **0.12**<br>(-.012 – 0.37) | **1.87\*\***<br>(0.93-2.80) | **1.12\*\***<br>(0.39-1.85) | **1.49\*\***<br>(0.66-2.33) | **4.61\*\***<br>(3.71-5.50) | 803 |

Exchange ranks for crossovers on chromosome 2L
|  | No CO | Single CO | Double CO | Triple CO | E<sub>0</sub> | E<sub>1</sub> | E<sub>2</sub> | E<sub>3</sub> |
| --- | --- | --- | --- | --- | --- | --- | --- | --- |
| **Oregon-RM** | 585 | 377 | 9 | 0 | .223 | .739 | .037 | 0 |
| ***mei-p26<sup>1</sup>*** | 1419 | 796 | 103 | 3 | .312 | .516 | .162 | .01 |

### CO patterning analyses

CO interference was calculated as 1-coefficient of coincidence (*coc*), where *coc* is the observed number of double COs/expected number of double COs. The expected number of double COs is calculated as (CO frequency in interval 1)*(CO frequency in interval 2)*(total sample size).

The centromere effect was calculated as in [20]. Briefly, the centromere effect value was calculated as 1-observed number of COs/expected number of COs in the centromere spanning or centromere adjacent interval. The expected number of COs was calculated as (the total number of COs along the chromosome)*(physical length of the interval/total length of the chromosome). This calculation gives an estimate of what the CO frequency should be if CO frequency was proportionate to physical distance.

CO assurance was calculated as in [20]. Using the equations of Weinstein, the number of observed parental, single, double, and triple COs were converted to “bivalent exchange ranks”. This conversion is necessary because within a pair of homologous chromosomes (a bivalent), not all chromatids are involved in a CO (exchange) event. This means that for every bivalent, there are some chromatids that did not have a CO (non-exhange), and these chromatids can be placed in the gametes. Thus, the parental class of offspring that did not have a CO represent non-exchange chromosomes within an exchange bivalent and true non-exchange bivalents where no CO occurred. Similar arguments can be made for single, double, and triple CO classes. The equations of Weinstein account for the need to convert CO frequencies to bivalent exchange rates. The expected number of non-exchange bivalents was calculated using a Poisson distribution as in [20]. A Fisher’s exact test was used to compare expected and observed frequencies.

## Results

### Germline cyst development is delayed in mei-P26^1^ germaria

The *mei-P26*^*1*^ allele is a hypomorphic mutation caused by a P-element insertion in the first intron of the gene [11]. Unlike *mei-P26* null alleles, which have tumorous ovaries and severe fertility defects, *mei-P26*^*1*^ was previously reported to exhibit mild tumor phenotypes and normal fertility [11]; however, the mitotic and differentiation phenotypes were not rigorously quantified. To understand the extent of mitotic defects in *mei-P26*^*1*^ germaria, we analyzed well-established markers of cyst development: the fusome protein Hu-li tai shao (Hts), whose spatial localization serves as an excellent indicator of cyst identity [21]; Bam-GFP, enriched in 2-, 4-, and 8-cell cysts [22]; and the oocyte-enriched proteins Orb [23] and C(3)G (a component of the synaptonemal complex; [15]). Wildtype germaria have a stereotypical pattern of cyst progression, wherein cysts divide and move posteriorly in the germarium. As a consequence of differing rates of mitotic division, germaria have variable number of cysts (Figure 1B). When cysts transition from mitosis to meiosis, they express Bam-GFP and Orb in non-overlapping stages of differentiation (n = 20 germaria, Figure 1C). In contrast, while the overall morphology of *mei-P26*^*1*^ germaria was largely preserved, the composition of cysts within these germaria differed significantly from wildtype (Genotype × Stage interaction: χ^2^(5) = 16.27, p = 0.006; Conway-Maxwell Poisson mixed model; see Methods for details; Figure 1B, D). Specifically, *mei-P26*^*1*^ germaria accumulated more 2-cell cysts (Mean Ratio = 1.76 ± 0.35, p = 0.005), 4-cell cysts (Mean Ratio 2.04 ± 0.51; p = 0.004), and 16-cell cysts (Mean Ratio = 1.54 ± 0.15; p < 0.001; Figure 1B,D). Although 24% of *mei-P26*^*1*^ germaria displayed mutually exclusive expression of Bam and Orb consistent with wildtype (Figure 1D, n = 29 germaria), most had one or more cysts that co-expressed Bam-GFP and Orb (62%, Figure 1E). Rarely, we also observed *mei-P26*^*1*^ germaria that expressed Bam but failed to up-regulate Orb (14%, Figure 1F). Consistently, the cell cycle regulator Cyclin A is stabilized by Bam such that it is robustly expressed in the anterior tip of wildtype germaria (roughly corresponding to the mitotically dividing region) and downregulated by region 2a after the last mitotic division (Supplemental Figure 1A) [24]. Yet in *mei-P26*^*1*^ germaria, we observed co-expression of Cyclin A and Orb in region 2b cysts in 13% of germaria (n = 32 germaria, Supplemental Figure 1B) far beyond the region where mitotic divisions are typically complete. Despite the abnormal timing of cyst division, a single Orb- and C(3)G-positive oocyte was specified by region 3 in all wildtype germaria (n = 44) and in 80% of *mei-P26*^*1*^ germaria (n = 49, p = 0.003, Fisher’s exact test, Figure 1C’-F’). Remarkably, *mei-P26*^*1*^ homozygous females are fertile despite the mitotic defects.

Together, these results show that unlike *mei-P26* null mutants that completely block germ cell differentiation prior to meiotic initiation [11, 13], the hypomorphic *mei-P26*^*1*^ mutant slows, but does not prevent, cyst division and differentiation, likely due to the complex interaction between Mei-P26 and Bam [12]. The co-expression of Bam and Orb in a subset of cysts leads to the intriguing possibility that some cysts in this genetic background receive mitotic signals while also initiating oocyte specification.

### The synaptonemal complex rarely fully assembles in mei-P26^1^

The moderate developmental phenotypes of the *mei-P26*^*1*^ mutant allowed us to examine how delayed differentiation affects meiotic chromosome dynamics. Previous studies reported impaired CO formation and high chromosome nondisjunction in *mei-P26*^*1*^ mutants [11], and hinted that the synaptonemal complex might not fully form [15]. To more fully understand the extent of meiotic progression in *mei-P26*^*1*^ mutants, we visualized synaptonemal complex assembly by immunostaining for the transverse filament C(3)G and categorized its morphology as foci, discontinuous, or continuous (Figure 2). In wildtype, the synaptonemal complex is loaded at centromeres in 8-cell cysts and then into patches on chromosome arms in 16-cell cysts. By the time the 16-cell cyst enters region 2a, full length continuous synaptonemal complex is loaded along chromosome arms. This process is reflected in our quantitative staging of synaptonemal complex assembly in wildtype: fewer than 10% of GSCs, cystoblasts, and 2-cell cysts contained C(3)G foci, increasing to 37% and 48% in 4- and 8-cell cysts (Figure 2I). By the 16-cell stage, 91% of wild-type cysts displayed continuous synaptonemal complexes (Figure 2I). We also measured C(3)G fluorescence intensity and see a dramatic increase in intensity beginning at the 8-cell stage and peaking at the 16-cell stage (Supplemental Figure 2).

**Figure 2.**
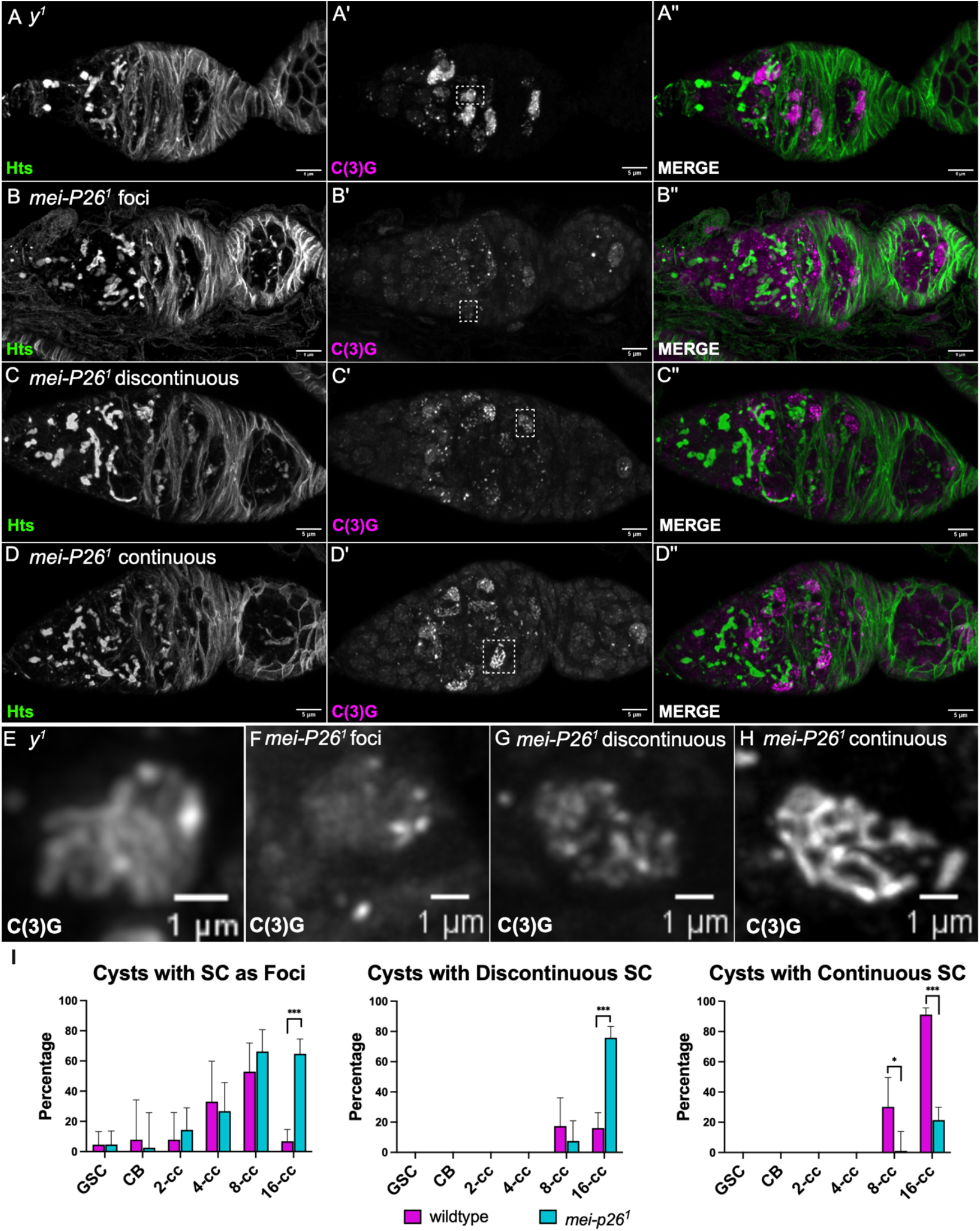
The synaptonemal complex does not fully assemble in *mei-P26*^*1*^. **A-A’’)** In *y*^*1*^ control germaria, C(3)G, the transverse filament of the synaptonemal complex, is loaded at centromeres during the mitotic divisions and can be seen as individual foci. In the 16-cell cyst, C(3)G is loaded into continuous tracks on chromosome arms in 2-4 cells within the cyst. As the 16-cell cyst moves posteriorly into region 2b, the synaptonemal complex is maintained in the two pro-oocytes and any other nuclei that initiated meiosis begin to degrade the synaptonemal complex. By region 3, only a single nucleus maintains the synaptonemal complex in the designated oocyte and the other pro-oocyte degrades the synaptonemal complex. A partially degraded synaptonemal complex can be seen adjacent to the oocyte in A’’. **B-B’’)** An example of a *mei-P26*^*1*^ germarium where the SC is never loaded onto chromosome arms and can only be seen as foci. **C-C’’)** An example of a *mei-P26*^*1*^ germarium where the synaptonemal complex is loaded into discontinuous tracks. **D-D’’)** An example of a *mei-P26*^*1*^ germarium where continuous synaptonemal complex is loaded onto chromosome arms. Note that not all meiotic nuclei load continuous synaptonemal complex in this germarium. **E-H)** Zoomed views of the nuclei in white dashed boxes showing various states of synaptonemal complex morphology. **I)** Quantification of synaptonemal complex morphology by cyst stage, *mei-P26*^*1*^ cysts frequently contained more than one type of morphology so percentages do not equal 100%, see text for statistical details, *p = 0.01, *** p < 0.001

In *mei-P26*^*1*^ germaria, C(3)G fluorescence intensity was modestly elevated in GSCs (MR = 1.16 ± 0.08, p = 0.036) and 4-cell cysts (1.21 ± 0.12, p = 0.041; Supplemental Figure 2), but this signal appeared diffuse throughout the nucleus rather than organized into synaptonemal complex (Figure 2I). By contrast, C(3)G intensity was reduced in 8-cell cysts (MR = 0.84 ± 0.07, p = 0.032) and 16-cell cysts (MR = 0.79 ± 0.05, p < 0.001; Supplemental Figure 2), coinciding with the stages at which synaptonemal complex normally matures from centromeric foci to full assembly. Continuous synaptonemal complex formation was severely compromised at the 16-cell cyst stage. Only 21.7% of *mei-P26*^*1*^16-cell cysts displayed continuous C(3)G compared to 89.8% in wildtype (Bayesian-penalized binomial mixed model; Figure 2; Supplementary Table S1). This reduction reflected a strong genotype effect (χ^2^(1) = 23.57, p < 0.001) rather than a stage-specific Genotype × Stage interaction, which was not significant for continuous C(3)G (χ^2^(1) = 0.0002, p = 0.99). Instead, *mei-P26*^*1*^16-cell cysts were enriched for incomplete synaptonemal complex morphologies: 74.7% exhibited discontinuous C(3)G compared with 17.4% in wildtype (Odds Ratio = 16.3 ± 6.63, p < 0.001; Stage × Genotype interaction: χ^2^(1) = 18.65, p < 0.001) and 63.6% displayed C(3)G foci compared with 8.1% in wildtype (Odds Ratio = 25.3 ± 12.70, p < 0.001; Stage × Genotype interaction: χ^2^(5) = 30.40, p < 0.001; Bayesian-penalized binomial mixed model). Co-staining with the centromere marker Cenp-C confirmed that C(3)G foci in both genotypes localize to centromeres, indicating that initial synaptonemal complex nucleation is intact (Figure 3). Together, these data suggest that the earliest stages of synaptonemal complex assembly in *mei-P26*^*1*^ occur normally, but *mei-P26*^*1*^ specifically often fails to fully assemble the synaptonemal complex in 16-cell cysts.

**Figure 3.**
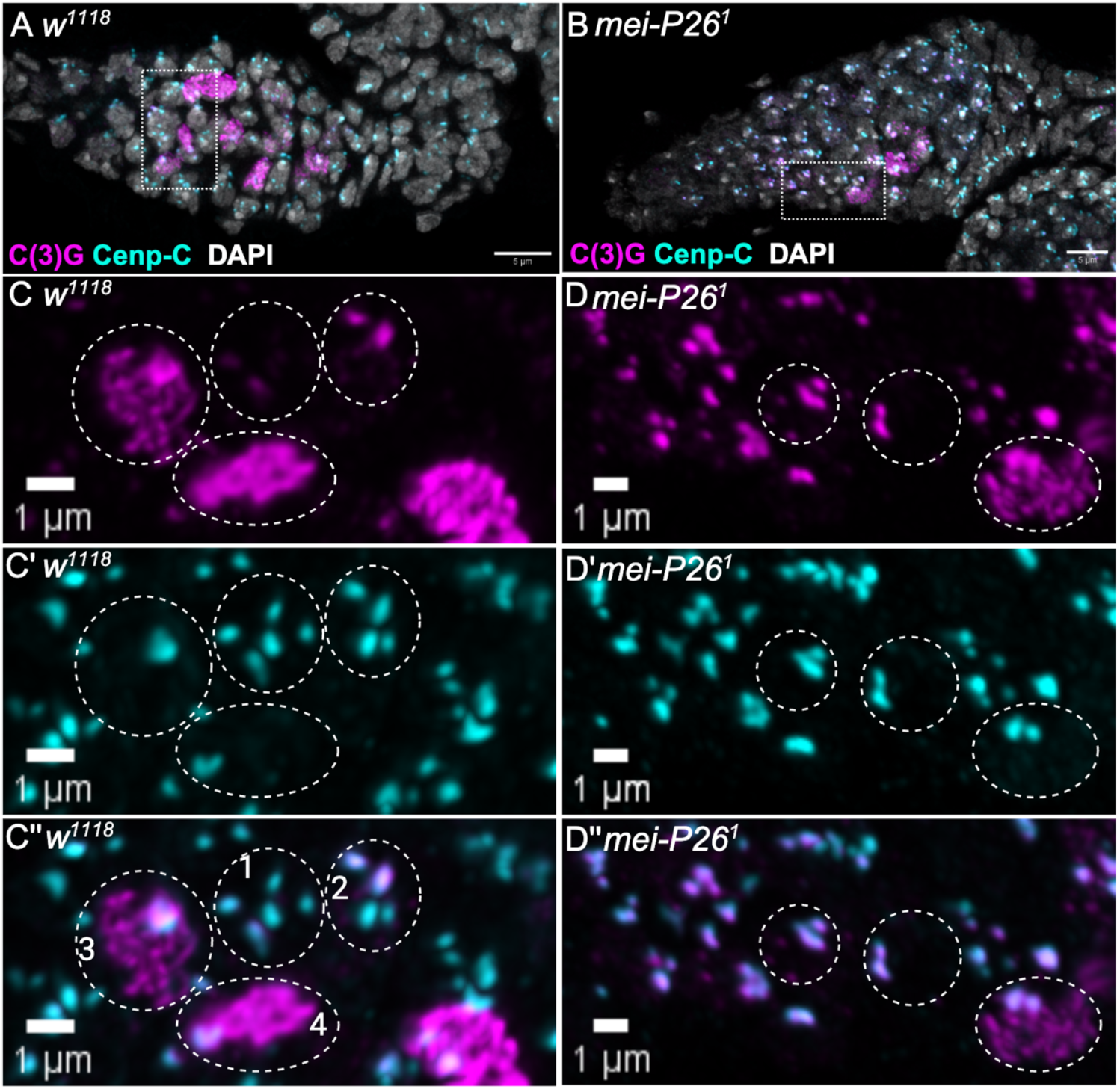
*mei-P26*^*1*^ loads centromeric synaptonemal complex. **A and B)** Germaria from *w*^*1118*^ controls and *mei-P26*^*1*^ immunostained with Cenp-C, C(3)G and DAPI. **C-C’’)** Zoomed in view of dashed box in A. Circles represent individual nuclei at various stages of centromere loading and arm loading of C(3)G. Nucleus 1 has just started to load C(3)G on one centromere. Nucleus 2 has C(3)G loaded onto two centromeres. Nucleus 3 has C(3)G fully loaded on centromeres, the centromeres are clustered, and loading of arm C(3)G has begun. Nucleus 4 has complete centromere and arm C(3)G. **D-D’’)** Zoomed in view of dashed box in B. Circles represent individual nuclei at various stages of centromere loading and arm loading of C(3)G. The two left nuclei have loaded centromeric C(3)G but no arm C(3)G. The bottom right nucleus has loaded centromere C(3)G and begun loading arm C(3)G.

We next asked whether defective synaptonemal complex assembly correlates with aberrant Bam–Orb expression. In control germaria, Bam-positive cysts lacked synaptonemal complex or showed only C(3)G foci, whereas Orb-positive cysts had fully assembled synaptonemal complex (Figure 1C). In *mei-P26*^*1*^, Bam-only cysts showed similarly limited synaptonemal complex assembly, and Orb-only cysts displayed discontinuous or continuous synaptonemal complex (Figure 1D-F). Strikingly, cysts co-expressing Bam and Orb lacked synaptonemal complex or contained only C(3)G foci (Figure 1E). These results suggest that inappropriate persistence of Bam interferes with C(3)G extension along chromosome arms, and that timely downregulation of Bam may be required for full synaptonemal complex assembly during meiosis.

### Meiotic double-strand breaks form in mei-P26^1^

In *Drosophila*, the majority of meiotic double-strand break (DSB) formation requires the presence of the synaptonemal complex; in *c(3)g* null mutants that fail to make a synaptonemal complex, DSB frequency is reduced by 80% [8, 9]. Because *mei-P26*^*1*^ forms COs [11] despite defects in synaptonemal complex assembly, we examined meiotic DSB formation by immunostaining against phosphorylated H2AV (γH2AV), a conserved histone modification made to initiate the DNA damage response to meiotic and somatic DSBs (Figure 4) [9, 25]. In wildtype, γH2AV appears at background levels or in cells with DNA damage in the GSCs, CB, 2-, 4-, and 8-cell cysts (Figure 4A-A’). After the synaptonemal complex is fully assembled in 16-cell cysts, γH2AV signal increases sharply (Figure 4E), indicating the induction of meiotic DSBs. These breaks are repaired by region 3 (Figure 4A-A’ and F).

**Figure 4.**
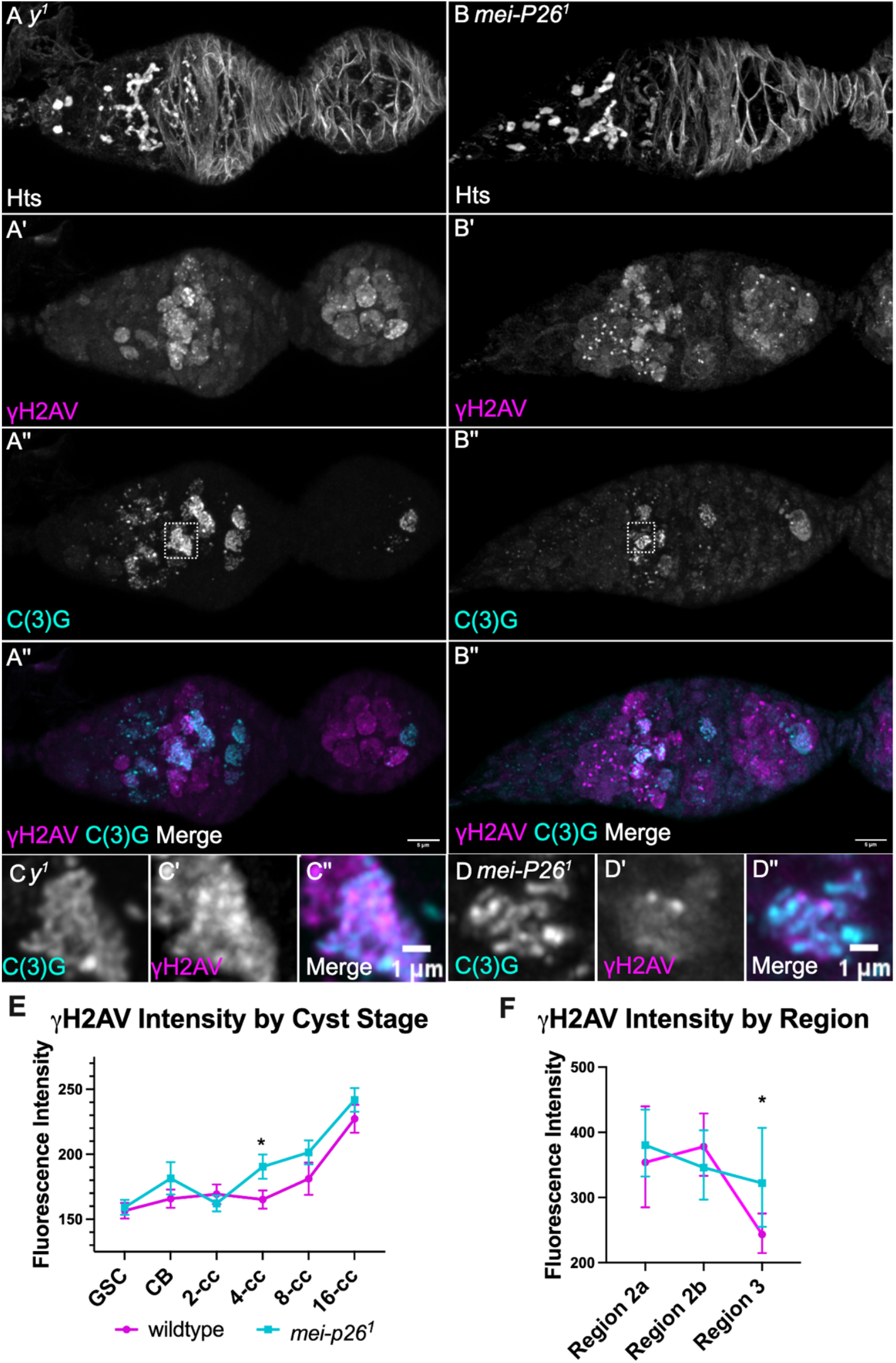
Meiotic double-strand breaks form at wildtype levels in *mei-P26*^*1*^. **A-A’’)** In control *y*^*1*^ germaria, there is a large induction of γH2AV signal in region 2a, corresponding to the formation of meiotic DSBs. DSBs are mostly repaired by region 2b, as can be seen by the lack of γH2AV. All DSBs should be repaired by region 3. **B-B’’)** In *mei-P26*^*1*^ germaria, γH2AV representing meiotic DSBs form. In this particular example, γH2AV can be seen in region 2a, but there is no signal in regions 2b and 3, indicating DSBs have been repaired. See text and Supplemental Figure 3 for discussion of the variability in DSB formation and repair. **C-D)** Zoomed in views of the dashed white boxes in A’’ and B’’ showing the pattern of γH2AV. **E)** γH2AV fluorescence intensity was measured in whole cysts for GSCs through 16-cell cysts. 16-cell cysts include those found in regions 2a, 2b and 3. **F)** γH2AV fluorescence that was present along the synaptonemal complex was measured in C(3)G positive nuclei from 16-cell cysts and grouped by region. For both E and F, *p = 0.027, see text for statistical methods.

In *mei-P26*^*1*^ germaria, we observed three distinct γH2AV patterns. Diffuse whole-nucleus signal appeared in non-meiotic nuclei lacking C(3)G randomly distributed throughout the germarium (Supplemental Figure 3A), or in clusters of nuclei with intense γH2AV signal, consistent with apoptosis (Supplemental Figure 3B). Both patterns differ from normal meiotic double-strand break timing and distribution and likely represent non-specific DNA damage.γH2AV was also observed in nuclei with either discontinuous or continuous synaptonemal complex (Figure 4B-D). To determine whether these signals represent meiotic DSBs or DNA damage, we generated *mei-P26*^*1*^; *mei-p22*^*P22*^ double mutants and analyzed H2AV phosphorylation [26, 27]. In these mutants, nuclei containing synaptonemal complex lacked γH2AV signal (Supplemental Figure 3C), demonstrating that the observed DSBs are *mei-p22*-dependent and therefore genuine meiotic breaks.

As a proxy for counting the number of DSBs formed in each genotype, we quantified γH2AV fluorescence intensity (see Methods for details). For GSCs through 8-cell cysts, we measured fluorescence across the entire cyst (Figure 4E); for 16-cell cysts, we quantified γH2AV fluorescence signal that co-localized with C(3)G (Figure 4F). Average fluorescence intensities from GSCs through 8-cell cysts did not differ significantly between wildtype and *mei-P26*^*1*^ germaria, except for a small increase in *mei-P26*^*1*^ 4-cell cysts (Mean Ratio = 1.15 ± 0.074, p = 0.027; heterogeneous-variance mixed model; Genotype × Stage interaction, χ^2^(5) = 12.78, p = 0.026; Figure 4E). In C(3)G-positive nuclei of 16-cell cysts, γH2AV intensity was similar in regions 2a and 2b but significantly higher in region 3 in *mei-P26*^*1*^ compared to wildtype, (Mean Ratio = 1.32 ± 0.18, p = 0.037; heterogeneous-variance mixed model; Genotype × Region interaction, χ^2^(2) = 7.23, p = 0.027; Figure 4F).

Mean γH2AV intensity (Figure 4) captures the average amount of DSB-associated signal but not how synchronized that signal appears across cysts within each region. We therefore examined cyst-to-cyst variability in γH2AV signal among 16-cell cysts in regions 2a, 2b, and 3. Dispersion was significantly elevated in *mei-P26*^*1*^ in region 2b (Mean Ratio = 1.64 ± 0.33, p = 0.014) and region 3 (Mean Ratio = 2.98 ± 1.3, p = 0.012; Genotype × Region interaction, χ^2^(2) = 12.9, p = 0.002), while region 2a was not significantly different (Mean Ratio = 0.765 ± 0.12, p = 0.084; Supplemental Figure 4;). Together with the elevated mean γH2AV in region 3, these findings suggest that *mei-P26*^*1*^ largely retains the capacity to form meiotic DSBs but disrupts the synchrony and timing of DSB repair in late pachytene.

### Crossover formation and patterning are severely disrupted in mei-P26^1^

Previous studies showed that total CO frequency in *mei-P26*^*1*^ is reduced to 67% of wildtype and is unevenly distributed: COs are reduced on the chromosome arm, but are increased near the centromere [11]. These results suggest that COs still form, but CO patterning mechanisms might be disrupted. To test this directly, we measured CO frequency and distribution on chromosome 2L and chromosome 3.

Although our *mei-P26*^*1*^ stock has the same rate of X chromosome nondisjunction as previously reported [wildtype nondisjunction is 0% (n= 901); *mei-P26*^*1*^ nondisjunction is 26.6% (n= 423)], we did not observe a statistically significantly difference in total CO frequency (40.68 cM in wildtype *vs*. 43.89 cM in *mei-p26*^1^, p=.10, Fishers’ Exact Test, Table 1 and Figure 5). However, consistent with previous studies, the CO distribution in *mei-P26*^*1*^ was significantly shifted. COs on the arm are reduced to 8-32% of wildtype levels, while COs in the centromere adjacent interval are increased to 171% of wildtype levels (*b – pr*, p < 0.0001 for all intervals, Fisher’s Exact Test). Most strikingly, COs are increased to 1600% of wildtype levels in the centromere-spanning interval (*pr – cn*, p < 0.0001, Fisher’s Exact Test). To ensure this phenotype was consistent across chromosomes, we also analyzed chromosome 3 (Supplemental Figure 5; Supplemental Table 1). Although total CO frequency was reduced to 51% of wild type, COs in the centromere-spanning interval increased by 404%, similar to chromosome 2L.

**Figure 5.**
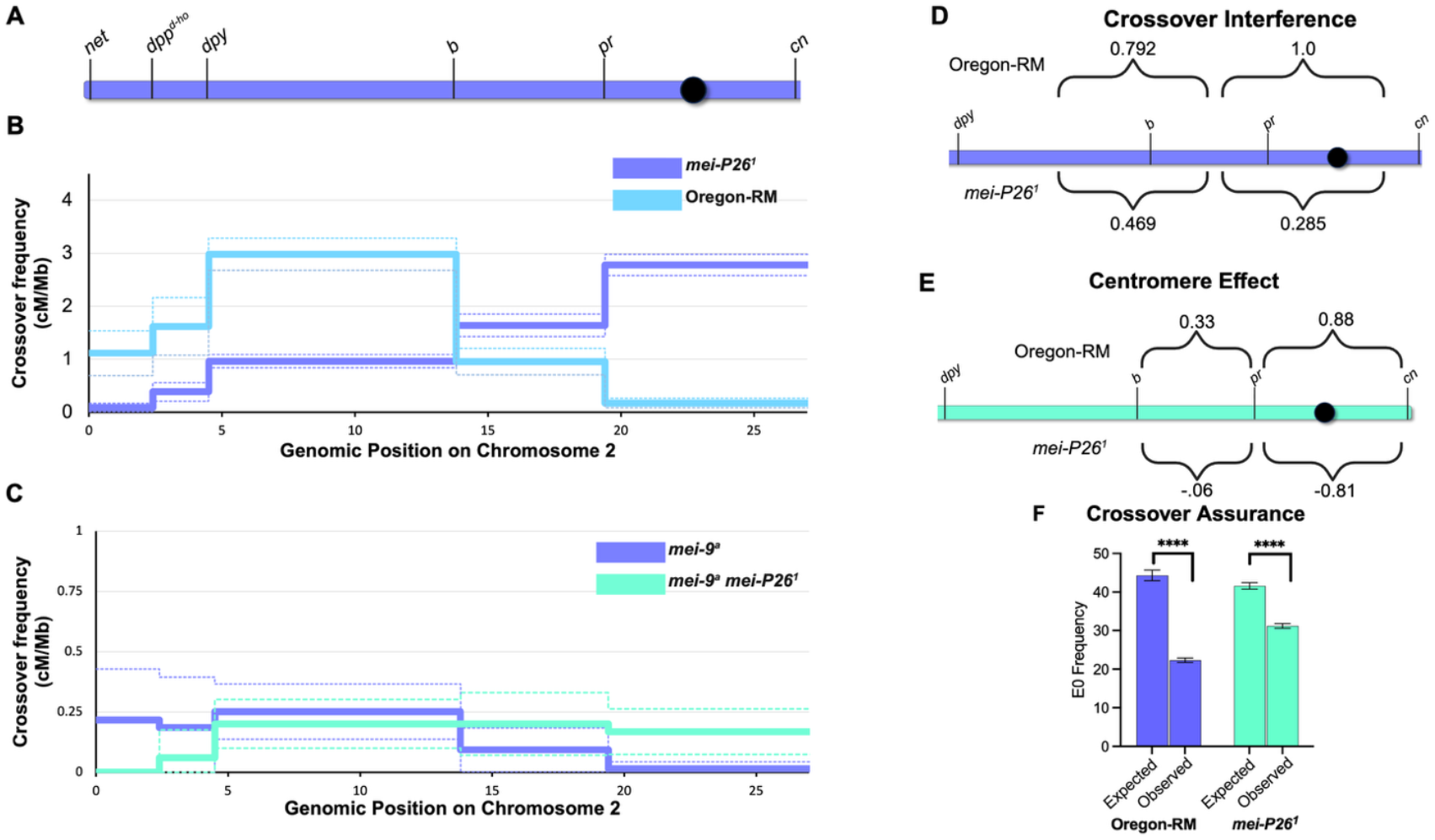
Crossover frequency and patterning in *mei-P26*^*1*^. **A)** Schematic showing the location of recessive markers used for scoring COs (drawn to scale). The black circle represents the centromere; chromosome 2R continues distal to *cn* but is not shown. **B)** CO frequencies in cM/Mb for Oregon-RM and *mei-P26*^*1*^. Dotted lines represent 95% confidence intervals. The total map length is the same for both genotypes, but COs are distributed differently in *mei-P26*^*1*^ compared to Oregon-RM. All intervals are statistically significantly different from Oregon-RM, see Table 1 for details. **C)** CO frequencies in *mei-9*^*a*^ and *mei-9*^*a*^ *mei-p26*^*1*^ double mutants. *mei-9* is required for 90% of COs. CO frequencies in *mei-9*^*a*^ *mei-p26*^*1*^ double mutants are not statistically different from *mei-9*^*a*^, showing that *mei-9* is required for the COs that form in *mei-p26*^*1*^. See Table 1 for details. Note the different scale of the y-axis. **D)** CO interference is significantly reduced in *mei-P26*^*1*^. **E)** The centromere effect is lost in *mei-P26*^*1*^. **F)** CO assurance is maintained in *mei-P26*^*1*^ because there are statistically significantly fewer chromosomes without a CO (E0) than expected by chance. See Methods for all details.

We next examined CO patterning mechanisms on chromosome 2L (interval spacing on chromosome 3 precluded these analyses). In wildtype flies, the presence of a CO in one genetic interval decreases the likelihood of a CO occurring in a neighboring genetic interval, and this CO interference keeps COs farther apart than expected by chance [28-30]. CO interference can be calculated as one minus the coefficient of coincidence (*coc*), where *coc* is the observed number of double COs divided by the expected number of double COs [31]. Interference values close to 1 mean CO interference is complete (such that one CO completely suppresses CO formation in the neighboring interval), values close to zero mean there is no CO interference, and negative values mean there is negative interference (such that a CO in that genetic interval increases the chance of a CO occurring the neighboring interval). Using intervals in which the CO frequencies were high enough to measure interference in both genotypes, we found interference in wildtype ranged from 0.792 to 1.0, showing that there is strong interference (Figure 5D). In *mei-P26*^*1*^, interference values decreased to 0.469 and 0.285, suggesting a decrease in CO interference (Figure 5D).

COs are normally suppressed in and near centromeres, a phenomenon known as the centromere effect [32-35]. The centromere effect can be quantitatively analyzed by comparing the observed number of COs to the expected number of COs if CO formation was proportionate to physical distance [20]. In this approach, values close to 1 represent suppression of COs, values close to zero represent no suppression of COs, and negative values represent promotion of COs. We found that in wildtype, the centromere effect value is 0.88 in the *pr-cn* interval but only 0.33 in the neighboring *b-pr* interval (Figure 5E). In *mei-P26*^*1*^, the centromere effect in *pr-cn* is -0.81 and -.06 in *b-pr*, showing that the centromere effect is lost in *mei-P26*^*1*^(Figure 5E). This is consistent with the large increase in CO frequency in the *pr-cn* interval (Figure 5B and Table 1).

Finally, we assessed CO assurance, which ensures that each homolog pair receives at least one CO to ensure accurate chromosome segregation. We measured CO assurance by calculating the number of chromosome pairs that did not receive a CO using the equations of Weinstein and comparing that to the expected number of chromosome pairs with no COs if CO formation was random across the genome (Methods and [20, 36]). In both wild type and *mei-P261*, the observed number of chromosomes lacking a CO was significantly lower than expected by chance, indicating that CO assurance remains intact (Figure 5F).

In Drosophila, mutations in *blm*, a master regulator of implementing meiosis-specific modes of homologous recombination, cause a flat distribution of COs, loss of CO assurance, loss of the centromere effect, loss of CO interference, and COs are no longer dependent on the putative Holliday junction resolvase Mei-9 [20]. Since some but not all CO patterning mechanisms are disrupted in *mei-P26*^*1*^, we asked whether COs require Mei-9 by generating a *mei-9*^*a*^ *mei-P26*^*1*^ double mutant and measuring CO frequencies on chromosome 2L. In *mei-9*^*a*^ single mutants, COs are reduced to 10% of wildtype levels (Table 1 and Figure 5C). COs are similarly reduced to 10% of wildtype levels in *mei-9*^*a*^ *mei-p26*^*1*^ double mutants (Table 1 and Figure 5C), showing that COs in *mei-p26*^*1*^ mutants require the canonical meiotic recombination machinery.

All together, these results show that in *mei-P26*^*1*^ total CO frequencies are approximately 50-100% of wildtype levels, but the distribution of those COs is significantly different, with a dramatic shift towards the centromere regions and away from the chromosome arms. Consistent with this, the centromere effect is lost, and the strength of CO interference is decreased. Surprisingly though, CO assurance is still active, and COs require the meiotic recombination machinery to form.

## Discussion

In the current work, we investigated how meiotic chromosome dynamics are coordinated with the developmental programming of germ cell differentiation and oocyte fate acquisition. Most *Drosophila* mutations that affect developmental programming result in tumorous ovaries or infertility, precluding an analysis of meiotic chromosome dynamics. By using a fertile hypomorphic allele of *mei-P26* with crossover defects, we were able to directly examine these processes. We found that in *mei-p26*^*1*^, many cysts co-express Bam and Orb, but *bam* expression is eventually lost and an oocyte is usually specified by region 3. Concurrently, we found that although most *mei-P26*^*1*^ germ cells initiate meiosis, synaptonemal complex assembly is often incomplete. Meiotic DSBs still form and are repaired into COs, but CO patterning is disrupted. Based on these results, we propose a model where the failure to degrade Bam (and potentially other mRNAs necessary for mitotic division) in a timely fashion causes cells to enter or progress through meiosis while still receiving mitotic signals. It is tempting to speculate that a clear cell cycle identity is necessary to execute normal meiotic chromosome dynamics, but further support for this model requires information on how meiotic cell identity is established and if it is dependent or independent of oocyte fate acquisition.

Why might the failure to cleanly exit mitosis disrupt meiotic chromosome dynamics? Traditional descriptions of the *Drosophila* germarium describe region 1 as being mitotic and regions 2a, 2b and 3 as meiotic. However, several pieces of evidence show that the earliest stages of meiosis begin in the mitotic cysts. Centromeric synaptonemal complex loading and centromere clustering are seen as early as 4-cell cysts and peak in the 8-cell cysts, and the dynamic nuclear rotations that facilitate robust meiotic chromosome pairing occurs in the 8-cell cysts [6, 37, 38]. These early meiotic events coincide with Bam expression. Bam is normally downregulated in 16-cell cysts when full length SC is assembled and DSBs form [39]. It is possible that the persistent expression of Bam in *mei-P26*^*1*^ may delay the transition from early meiotic prophase (homolog pairing) to the later stages of prophase when recombination occurs. Supporting this idea, the synaptonemal complex in *mei-P26*^*1*^ appears as discontinuous or continuous only in Orb-positive cysts, whereas cysts co-expressing Bam and Orb have minimal synaptonemal complex assembly (Figure 1).

It is unclear why a hypomorphic mutation in *mei-P26* would lead to co-expression of Bam and Orb. One possible molecular pathway is that Mei-P26 is required for Rbfox (an RNA binding protein) expression [40], and Rbfox expression is required for degradation of Bam transcripts [41]. In this case, decreased *mei-P26* expression would lead to decreased *Rbfox* expression, which could prevent the timely degradation of *bam* transcripts.

Despite extensive disruption of CO patterning, *mei-P26*^*1*^ still requires Mei-9 for CO formation and retains CO assurance. This contrasts with *blm* mutants, where COs form at ∼70% of wild-type levels but all CO patterning, including assurance, is lost and COs no longer require Mei-9. CO assurance may therefore be linked to the activity of meiosis-specific resolvases such as Mei-9, which are thought to generate exclusively COs [42-44]. Surprisingly, X-chromosome nondisjunction remains high (26%) in *mei-P26*^*1*^ despite intact CO assurance. One explanation is that the peri-centromeric COs (which form at a high frequency in *mei-P26*^*1*^) satisfy the obligate CO requirement but fail to support proper segregation, due to their abnormal position. This phenotype may arise because *mei-P26*^*1*^ retains the ability to nucleate synaptonemal complex at centromeres but inefficiently extend continuous synaptonemal complex.

By using a hypomorphic allele of *mei-P26*^*1*^, we’ve demonstrated that not only is germ cell developmental programming important for entry into meiosis, it’s also critical for regulating chromosome dynamics during meiosis. Studying germ cell development or meiotic chromosome dynamics as isolated processes prevents us from having a comprehensive understanding of how germ cells propagate genetic material to the next generation; future work must consider how development and chromosome dynamics are integrated.

## Data availability

All data and stocks are available upon request. The authors affirm that all data necessary for confirming the conclusions of the article are present within the article, figures, and tables.

## Acknowledgements

We would like to thank members of the Crown and Ables labs and our anonymous reviewers for helpful comments on the manuscript. We would also like to thank the Developmental Studies Hybridoma Bank (created by the NICHD of the NIH and maintained at The University of Iowa, Department of Biology, Iowa City, IA 52242). Stocks obtained from the Bloomington Drosophila Stock Center (NIH P40OD018537) were used in this study.

## References

1. Hinnant, T.D., J.A. Merkle, and E.T. Ables, Coordinating Proliferation, Polarity, and Cell Fate in the Drosophila Female Germline. Frontiers in Cell and Developmental Biology, 2020. 8.

2. Hassold, T. and P. Hunt, To err (meiotically) is human: the genesis of human aneuploidy. Nature Reviews Genetics, 2001. 2(4): p. 280–291.

3. Pazhayam, N.M., C.A. Turcotte, and J. Sekelsky, Meiotic Crossover Patterning. Frontiers in Cell and Developmental Biology, 2021. 9.

4. Hughes, S.E., et al., Female Meiosis: Synapsis, Recombination, and Segregation in Drosophila melanogaster. Genetics, 2018. 208(3): p. 875–908.

5. Takeo, S., et al., Synaptonemal Complex-Dependent Centromeric Clustering and the Initiation of Synapsis in Drosophila Oocytes. Current Biology, 2011. 21(21): p. 1845–1851.

6. Christophorou, N., T. Rubin, and J.-R. Huynh, Synaptonemal Complex Components Promote Centromere Pairing in Pre-meiotic Germ Cells. PLOS Genetics, 2013. 9(12): p. e1004012.

7. Tanneti, Nikhila S., et al., A Pathway for Synapsis Initiation during Zygotene in Drosophila Oocytes. Current Biology, 2011. 21(21): p. 1852–1857.

8. Jang, J.K., et al., Relationship of DNA double-strand breaks to synapsis in Drosophila. Journal of Cell Science, 2003. 116(15): p. 3069–3077.

9. Mehrotra, S. and K.S. McKim, Temporal Analysis of Meiotic DNA Double-Strand Break Formation and Repair in Drosophila Females. PLOS Genetics, 2006. 2(11): p. e200.

10. Lake, C.M., et al., Vilya, a component of the recombination nodule, is required for meiotic double-strand break formation in Drosophila. eLife, 2015. 4: p. e08287.

11. Page, S.L., et al., Genetic Studies of mei-P26 Reveal a Link Between the Processes That Control Germ Cell Proliferation in Both Sexes and Those That Control Meiotic Exchange in Drosophila. Genetics, 2000. 155(4): p. 1757–1772.

12. Li, Y., et al., Mei-P26 Cooperates with Bam, Bgcn and Sxl to Promote Early Germline Development in the Drosophila Ovary. PLOS ONE, 2013. 8(3): p. e58301.

13. Neumüller, R.A., et al., Mei-P26 regulates microRNAs and cell growth in the Drosophila ovarian stem cell lineage. Nature, 2008. 454(7201): p. 241–245.

14. Gui, J., et al., Simultaneous activation of Tor and suppression of ribosome biogenesis by TRIM-NHL proteins promotes terminal differentiation. Cell Reports, 2023. 42(3): p. 112181.

15. Page, S.L. and R.S. Hawley, c(3)G encodes a Drosophila synaptonemal complex protein. Genes Dev, 2001. 15(23): p. 3130–43.

16. Lake, C.M., et al., The Development of a Monoclonal Antibody Recognizing the Drosophila melanogaster Phosphorylated Histone H2A Variant (γ-H2AV). G3 Genes|Genomes|Genetics, 2013. 3(9): p. 1539–1543.

17. Anderson, L.K., et al., Juxtaposition of C(2)M and the transverse filament protein C(3)G within the central region of <i>Drosophila</i> synaptonemal complex. Proceedings of the National Academy of Sciences, 2005. 102(12): p. 4482–4487.

18. Fellmeth, J.E., et al., A dynamic population of prophase CENP-C is required for meiotic chromosome segregation. PLOS Genetics, 2023. 19(11): p. e1011066.

19. Gelman, A., et al., A weakly informative default prior distribution for logistic and other regression models. The Annals of Applied Statistics, 2008. 2(4): p. 1360–1383, 24.

20. Hatkevich, T., et al., Bloom Syndrome Helicase Promotes Meiotic Crossover Patterning and Homolog Disjunction. Current Biology, 2017. 27(1): p. 96–102.

21. Cuevas, M.d. and A.C. Spradling, Morphogenesis of the Drosophila fusome and its implications for oocyte specification. Development, 1998. 125(15): p. 2781–2789.

22. Beachum, A.N., et al., β-importin Tnpo-SR promotes germline stem cell maintenance and oocyte differentiation in female Drosophila. Dev Biol, 2023. 494: p. 1–12.

23. Lantz, V., et al., The Drosophila orb RNA-binding protein is required for the formation of the egg chamber and establishment of polarity. Genes & Development, 1994. 8(5): p. 598–613.

24. Sugimura, I. and M.A. Lilly, Bruno Inhibits the Expression of Mitotic Cyclins during the Prophase I Meiotic Arrest of Drosophila Oocytes. Developmental Cell, 2006. 10(1): p. 127–135.

25. Rogakou, E.P., et al., DNA Double-stranded Breaks Induce Histone H2AX Phosphorylation on Serine 139*. Journal of Biological Chemistry, 1998. 273(10): p. 5858–5868.

26. Liu, H., et al., mei-P22 Encodes a Chromosome-Associated Protein Required for the Initiation of Meiotic Recombination in Drosophila melanogaster. Genetics, 2002. 162(1): p. 245–258.

27. Robert, T., et al., The TopoVIB-Like protein family is required for meiotic DNA double-strand break formation. Science, 2016. 351(6276): p. 943–949.

28. Sturtevant, A.H., The linear arrangement of six sex-linked factors in Drosophila, as shown by their mode of association. Journal of Experimental Zoology, 1913. 14(1): p. 43–59.

29. Berchowitz, L.E. and G.P. Copenhaver, Genetic Interference: Don’t Stand So Close to Me. CURRENT GENOMICS, 2010. 11(2): p. 91–102.

30. von Diezmann, L. and O. Rog, Let’s get physical - mechanisms of crossover interference. Journal of cell science, 2021. 134(10): p. jcs255745.

31. Stevens, W.L., The analysis of interference. Journal of Genetics, 1936. 32(1): p. 51–64.

32. Painter, T.S., A New Method for the Study of Chromosome Rearrangements and the Plotting of Chromosome Maps. Science, 1933. 78(2034): p. 585–586.

33. Dobzhansky, T., TRANSLOCATIONS INVOLVING THE THIRD AND THE FOURTH CHROMOSOMES OF DROSOPHILA MELANOGASTER. Genetics, 1930. 15(4): p. 347–399.

34. Dobzhansky, T., TRANSLOCATIONS INVOLVING THE SECOND AND THE FOURTH CHROMOSOMES OF DROSOPHILA MELANOGASTER. Genetics, 1931. 16(6): p. 629–658.

35. Hartmann, M., J. Umbanhowar, and J. Sekelsky, Centromere-Proximal Meiotic Crossovers in Drosophila melanogaster Are Suppressed by Both Highly Repetitive Heterochromatin and Proximity to the Centromere. Genetics, 2019. 213(1): p. 113–125.

36. Weinstein, A., COINCIDENCE OF CROSSING OVER IN DROSOPHILA MELANOGASTER (AMPELOPHILA). Genetics, 1918. 3(2): p. 135–172.

37. Christophorou, N., et al., Microtubule-driven nuclear rotations promote meiotic chromosome dynamics. Nature Cell Biology, 2015. 17(11): p. 1388–1400.

38. Cahoon, C.K. and R.S. Hawley, Flies Get a Head Start on Meiosis. PLOS Genetics, 2013. 9(12): p. e1004051.

39. McKearin, D.M. and A.C. Spradling, bag-of-marbles: a Drosophila gene required to initiate both male and female gametogenesis. Genes & Development, 1990. 4(12b): p. 2242–2251.

40. Tastan, O.Y., et al., Drosophila ataxin 2-binding protein 1 marks an intermediate step in the molecular differentiation of female germline cysts. Development, 2010. 137(19): p. 3167–76.

41. Samuels, T.J., et al., Destabilisation of bam transcripts terminates the mitotic phase of Drosophila female germline differentiation. Development, 2025. 152(5): p. DEV204405.

42. Zakharyevich, K., et al., Delineation of Joint Molecule Resolution Pathways in Meiosis Identifies a Crossover-Specific Resolvase. Cell, 2012. 149(2): p. 334–347.

43. De Muyt, A., et al., BLM helicase ortholog Sgs1 is a central regulator of meiotic recombination intermediate metabolism. Mol Cell, 2012. 46(1): p. 43–53.

44. Klein, Hannah L. and Lorraine S. Symington, Sgs1—The Maestro of Recombination. Cell, 2012. 149(2): p. 257–259.

