## Supplemental Figures for "Analysis of a hypomorphic *mei-P26* mutation reveals coordination between developmental programming of germ cells and meiotic chromosome dynamics"

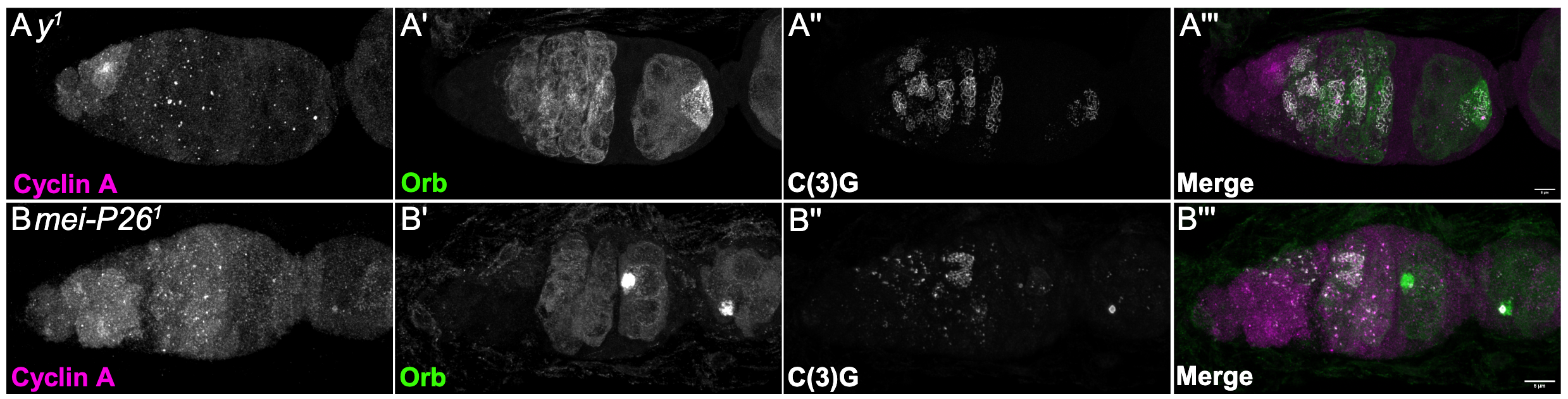
**Supplemental Figure 1.** Cyclin A expression is extended into region 2b in mei-P26^1^. **A)** In y^1^ controls, Cyclin A can be seen in a very early mitotically dividing cyst in the anterior end of the germarium. B) In mei-P26^1^, Cyclin A expression is extended, in this case, into region 2b (determined by morphology).

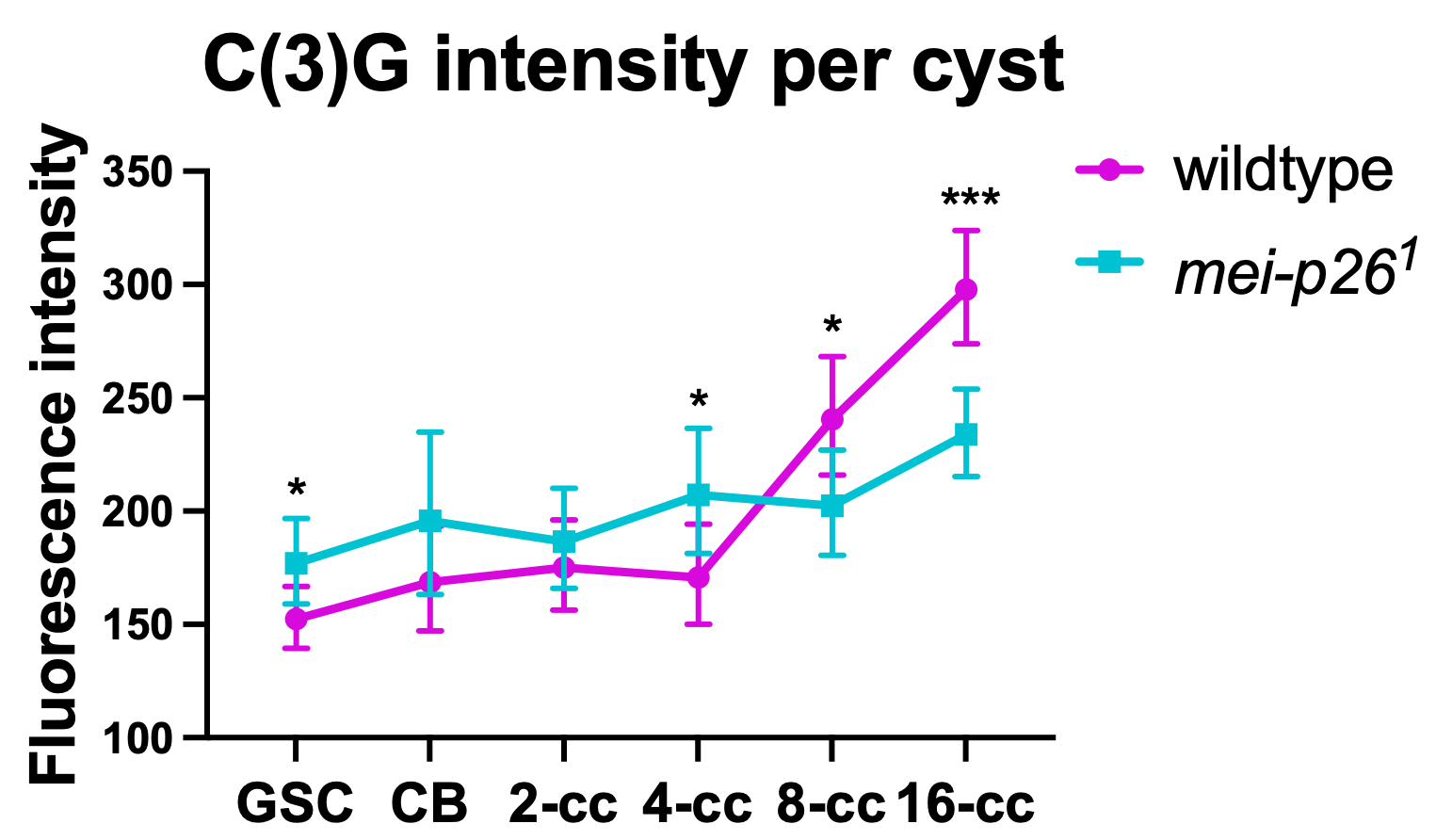

**Supplemental Figure 2.** C(3)G mean fluorescence intensity per cyst. 16-cell cysts include those found in regions 2a, 2b and 3. There is a dramatic induction of C(3)G expression beginning at the 8-cell stage. mei-P26^1^ germaria show statistically significantly higher C(3)G expression in GSCs and 4-cell cysts but significantly lower expression in the 8-cell and 16-cell cysts. See main text for statistical methods. See main text for statistical details; *p < 0.05, ***p < 0.001

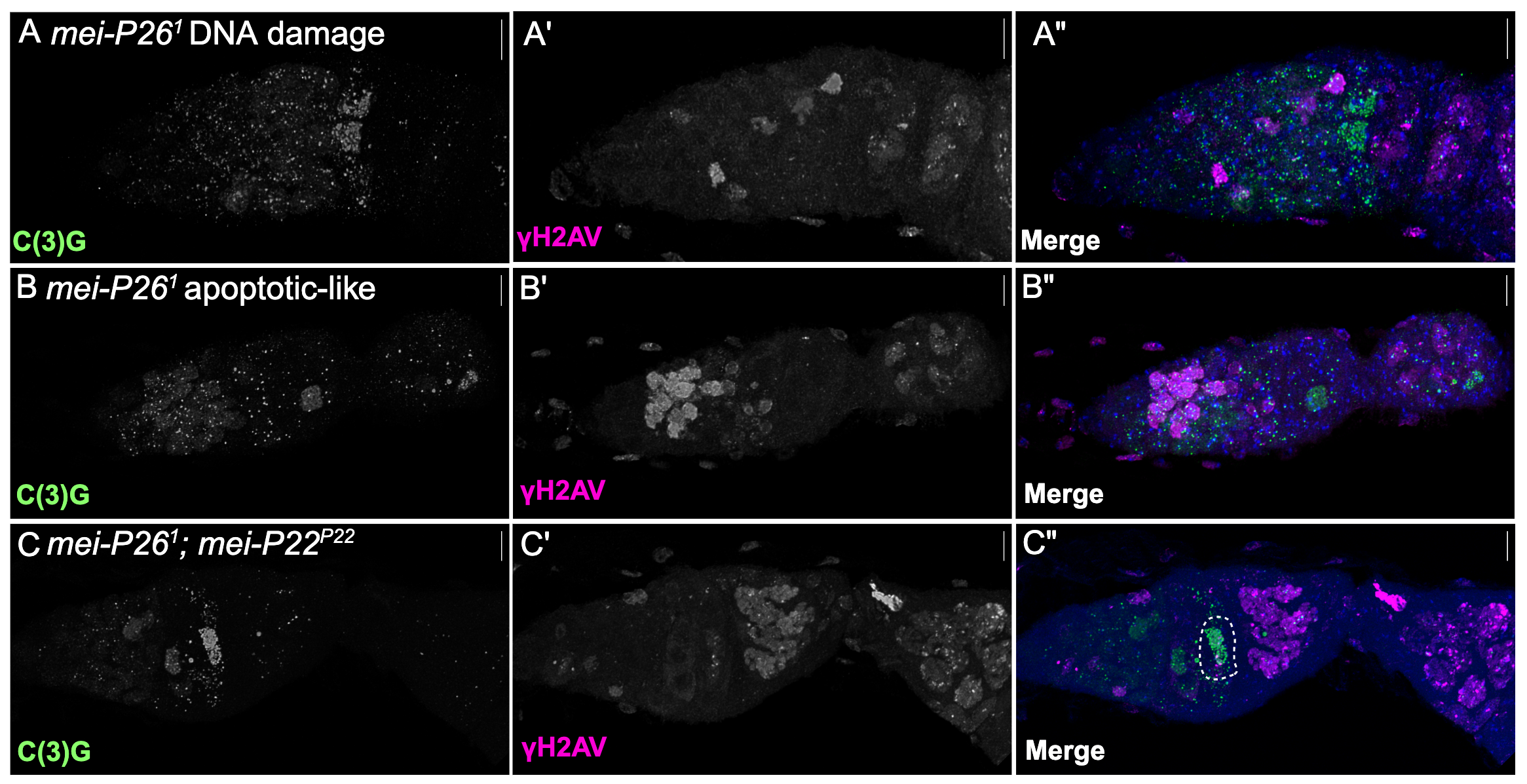

**Supplemental Figure 3.** γH2AV signal in mei-P26^1^. A) In some mei-P26^1^germaria, γH2AV is only detected as diffuse whole nucleus staining, likely representing background DNA damage. B) In some mei-P26^1^germaria, γH2AV signal is intense and appears clustered in one area, possibly representing a cyst undergo apoptosis. C) In mei-P26^1^; mei-P22^p22^ double mutants, only background levels of γH2AV are seen, suggesting that γH2AV that overlaps with C(3)G represents true meiotic double-stranded breaks.

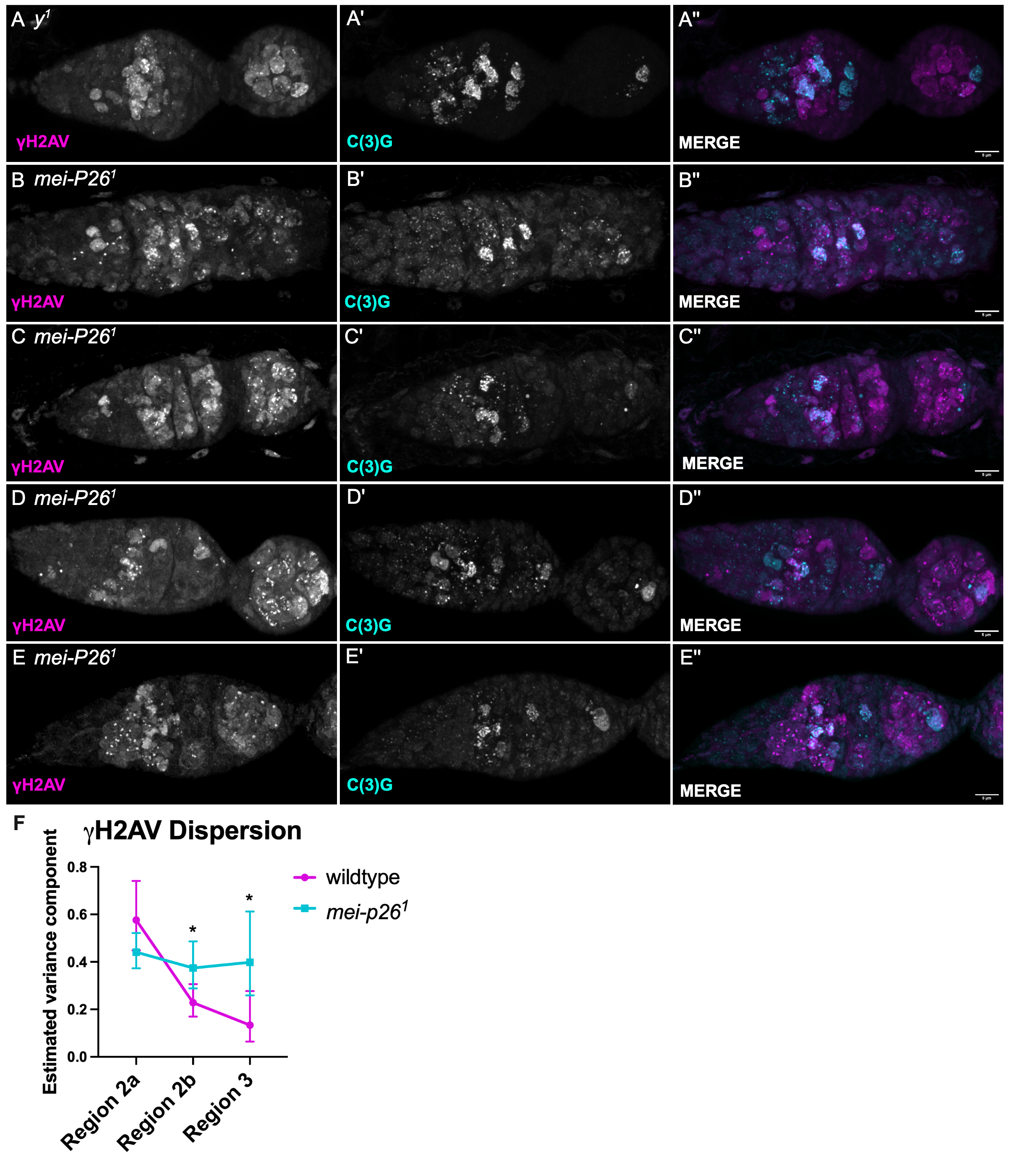

**Supplemental Figure 4. γH2AV signal in mei-P26^1^shows a variety of timing and location. A and E** are the same germaria shown in Figure 7 and are presented here for comparison. **B)** A mei-P26^1^ germarium with three region 2b cysts (based on morphology) with unrepaired DSBs. **C-D)** mei-P26^1^ germaria with DSBs in C(3)G positive nuclei. Cysts that are in region 2b (based on morphology) have repaired DSBs. **E)** A mei-P26^1^ germarium that has normal timing of DSB formation and repair (similar to y^1^), except there are fewer C(3)G positive nuclei than expected. There is additional γH2AV signal in an anterior cyst that might be in the early stages of apoptosis. **F)** See main text for statistical details; ***** p = 0.01

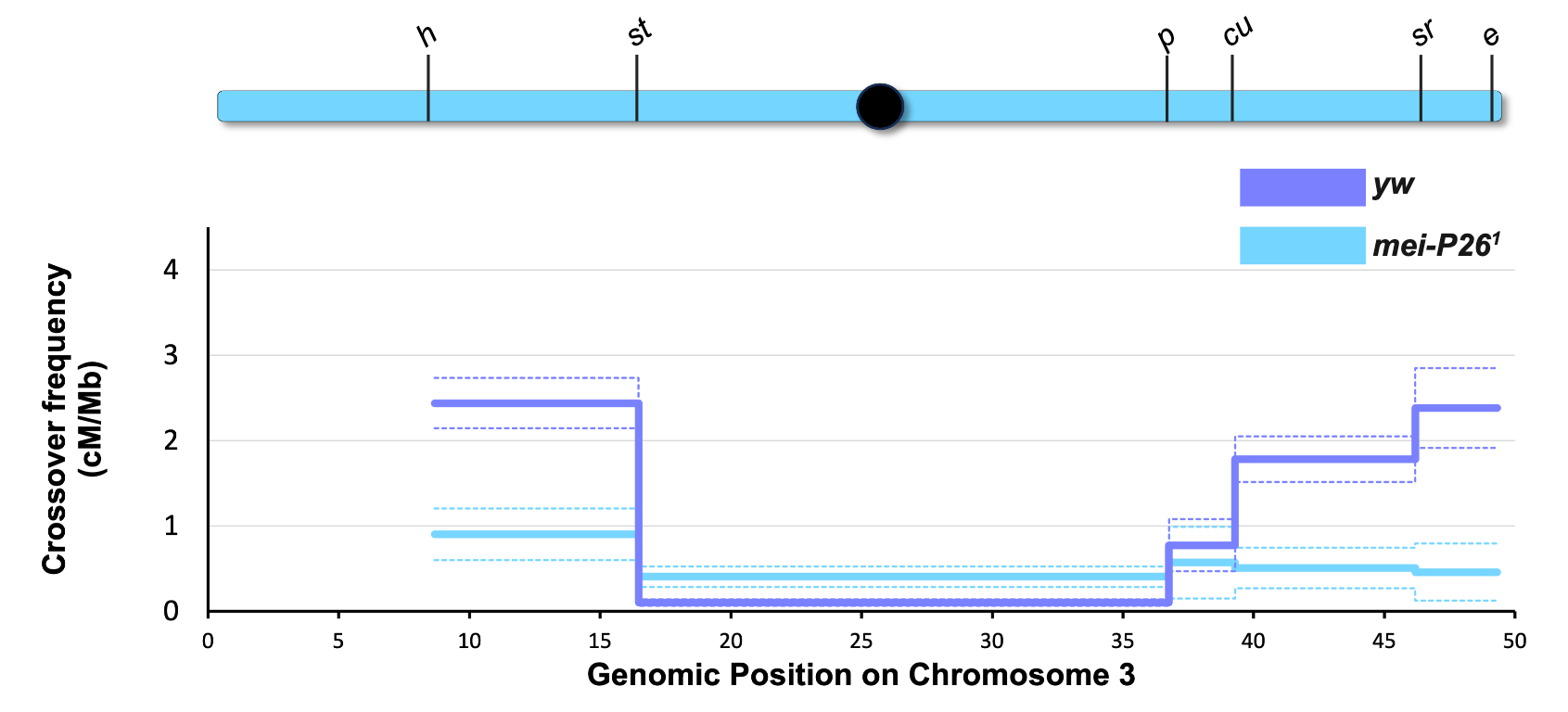

**Supplemental Figure 5.** Schematic showing the location of recessive markers used for scoring COs on chromosome 3 (drawn to scale). The black circle represents the centromere. CO frequencies in cM/Mb for yw and mei-P26^1^. Dotted lines represent 95% confidence intervals.

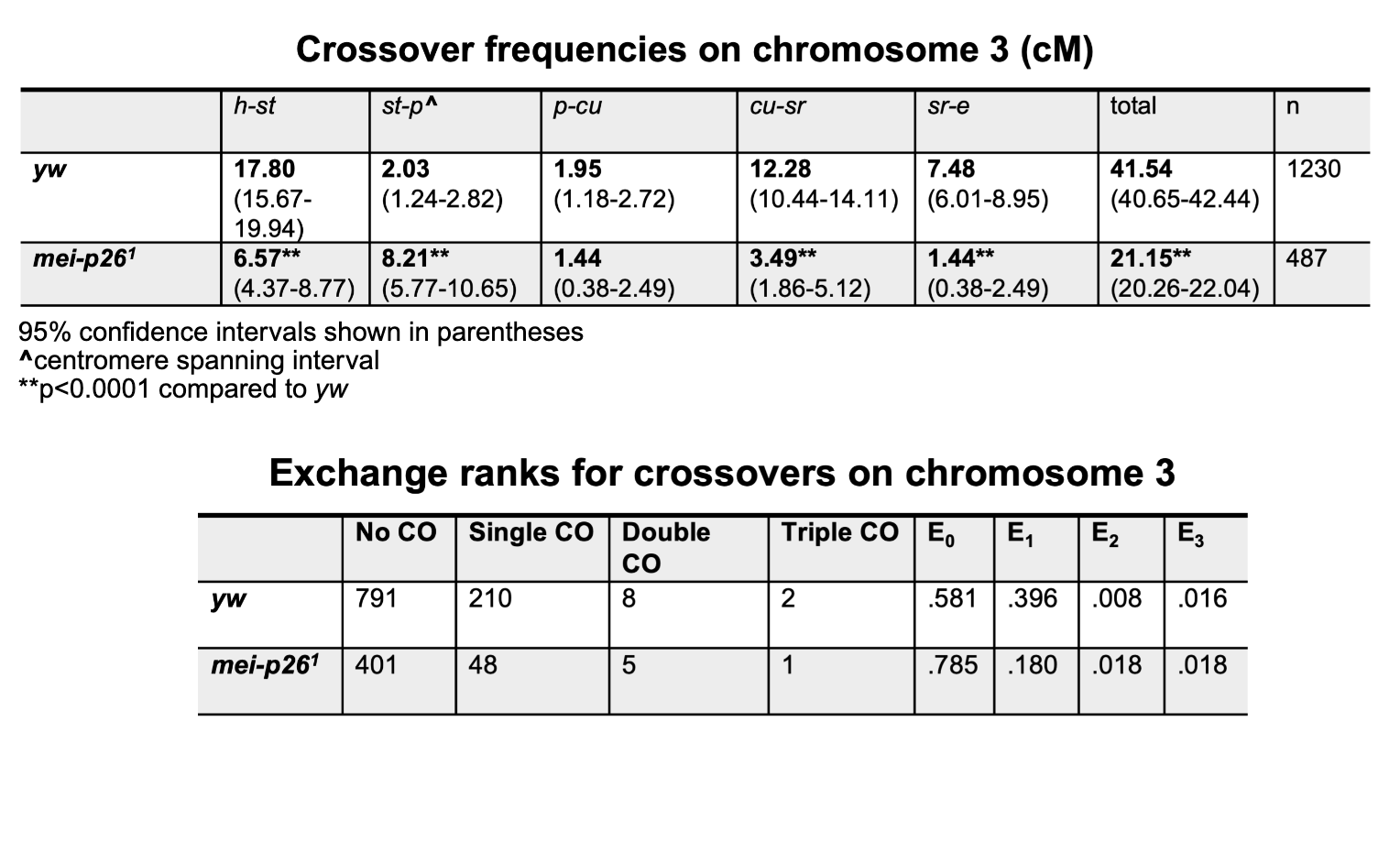

**Supplemental Table 1**
